# Molecular interaction and transport limitation in macromolecular binding to surfaces

**DOI:** 10.1101/380337

**Authors:** Pieter Wilhelm Hemker, Adam Miszta, Sonya M. Bierbower, Hendrik Coenraad Hemker, Wim Hermens

## Abstract

Binding of macromolecules to surfaces, or to surface-attached binding partners, is usually described by the classical Langmuir model, which does not include interaction between incoming and adsorbed molecules or between adsorbed molecules.

The present study introduces the “Surfint” model, including such interactions. Instead of the exponential binding behaviour of the Langmuir model, the Surfint model has tanh binding equations, as confirmed by a random sequential adsorption (RSA) computer simulation.

For high binding affinity, sorption kinetics become diffusion-limited as described by the existing unstirred-layer model “Unstir”, for which we present the exact analytical solution of its binding equations expressed in Lambert W-functions.

Low-affinity binding of thrombin on heparin, and high-affinity binding of prothrombin on phospholipid vesicles, were measured by ellipsometry and were best described by the Surfint and Unstir models, respectively.

## INTRODUCTION

Macromolecular binding on (bio)surfaces is important in biology, biocompatibility testing of implants, affinity screening of antibodies, DNA microarray analysis, testing anti-fouling agents, production of biosensors, etc. Such binding studies have been facilitated by solid-phase techniques like ellipsometry, surface plasmon resonance (SPR), quartz crystal microbalance (QCM) and total internal refection fluorescence (TIRF), allowing accurate real-time measurement of binding kinetics.

In the analysis of such studies, the classical Langmuir binding model [1] is still routinely used [2,3]. As apparent from Langmuir’s paper title: “Adsorption of gases on glass, mica and platinum”, he studied adsorption of small molecules on large matrix atoms and thus did not include interaction between incoming and adsorbed molecules or between adsorbed molecules. The model thus assumes constant values for the sorption constants kon and koff.

However, macromolecules are usually larger than their binding sites, as in the present study. Thrombin binds to small suger residues in the heparin chain [4], and prothrombin binds to small phosphatidylserine molecules in phospholipid membranes [5]. These proteins extend far beyond the binding sites, making binding dependent on adsorbed surface mass.

Indeed, deviations from Langmuirian adsorption have often been found and were related to electrostatic interactions, lateral mobility, cooperative binding, adsorption-induced molecular modification, surface reactions, etc. [6-17]. The many parameters involved in these interactions cannot be obtained from simple sorption curves, but in the present study they are lumped into a direct effect of adsorbed protein mass on sorption kinetics in a new “surface interaction” or Surfint model, applied to the low-affinity binding of thrombin on heparin.

For high-affinity binding, the fluid layer at the adsorbing surface will become depleted of adsorbent, making sorption kinetics diffusion-dependent. This is described by the existing unstirred-layer model ‘Unstir’, for which we present its exact analytical solution, and its application to the adsorption of prothrombin on surface-supported phospholipid vesicles.

## Materials and Methods

### The 4-parameter Langmuir binding model

As explained in the Introduction, this model assumes binding of non-interacting molecules on separated binding sites of equal affinity, which implies strict monolayer adsorption [1]. Binding is the net result of adsorption and desorption as obtained from the law of mass action:

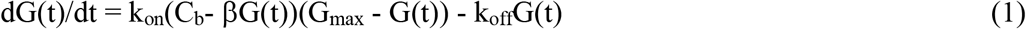

with G(t) the time-dependent surface concentration of sorbent, k_on_, k_off_ the rate constants of adsorption and desorption, C_b_ the initial bulk concentration of sorbent, βG(t) the lowering of C_b_ due to adsorption, and G_max_ the maximal surface concentration of sorbent, reached when all binding sites are saturated. Analytical solutions of this model have been presented in the literature for flat as well as spherical adsorbing surfaces [2, 3, 15]. In the present study only small amounts of protein were adsorbed and βG(t) could be neglected:

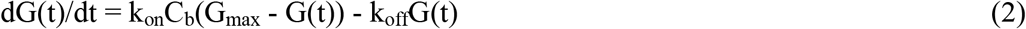

Starting adsorption at t = 0 and desorption (by reducing C_b_ to zero) at t = t_off_, we obtain the exponential sorption behaviour of the 4-parameter (C_b_, k_on_, k_off_, G_max_) Langmuir model:

<u>Adsorption, with G(t=0) = 0 and for t ≥ 0 a bulk concentration C_b_ of adsorbing protein:</u>

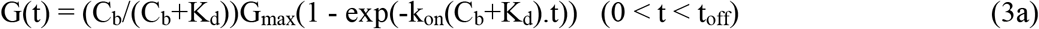

<u>Desorption, with G(t=t_off_) = G_off_ and C_b_= 0 for t ≥ t_off_:</u>

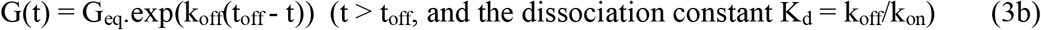

From (3a) we find that, for t → ∞, G(t) will reach its equilibrium value G_eq_:

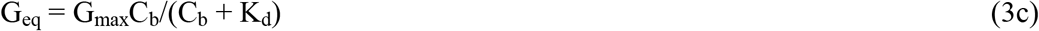

Values of G_max_ and K_d_ are usually obtained by fitting equation (3c) to G_eq_ values as obtained for a number of increasing C_b_ values.

### The 5-parameter unstirred-layer binding model (Unstir)

To limit adsorption time, protein binding is studied in stirred systems or in flow cells, and binding kinetics will depend on convection and diffusion. Directly at the surface, viscosity is dominant and flow velocity becomes zero - the “no slip” condition [18] - as used in the 5-parameter Unstir model [19], which assumes a thin layer of stagnant fluid at the surface, of thickness d, through which sorbents pass by diffusion. Beyond this layer one assumes a uniform bulk concentration C_b_ of sorbent, obtained by efficient stirring. Writing C(x,t) for the protein concentration at a distance x from the surface at time t, we obtain from the Langmuir model and the law of mass action:

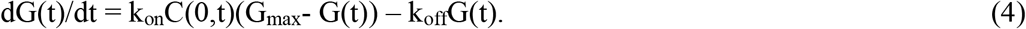

During sorption, the amount of protein in the unstirred layer is very small compared to the amount passing through it, causing rapid establishment of a quasi-steady state with *∂*C(x,t)/*∂*t = 0. From the diffusion equation *∂*C(x,t)/*∂*t = D *∂*C(x,t)/*∂*x^2^, with D the diffusion constant, it then follows that a linear gradient of sorbent will be established in the boundary layer within a second [20] and we obtain:

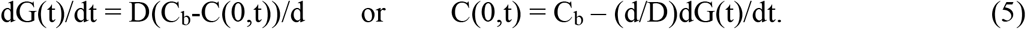

Substituting (5) into (4) we obtain after rearrangement of terms the Unstir binding equation:

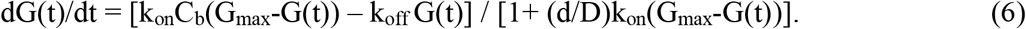

Equation (6) is identical to equation (2) with k_on_ and k_off_ replaced by respectively k_on,app_= k_on_/[1+(d/D)k_on_(G_max_-G(t))] and k_off,app_= k_off_/[1+(d/D)k_on_(G_max_-G(t))]. These constants thus become time-dependent but still allow determination of the dissociation constant K_d_ = k_off,app_/k_on,app_ = k_off_/k_on_. In the literature, equation (6) has only been solved numerically or as an infinite series [21], but we found that this equation allows for an exact analytical solution, presented below.

<u>Adsorption, with G(t=0) = 0 and for t ≥ 0 a bulk concentration C_b_ of adsorbing protein:</u>

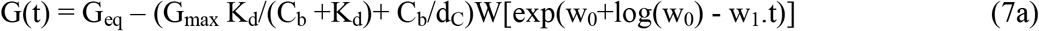

with G_eq_= G_max_C_b_/(C_b_+K_d_), d_C_ = k_on_C_b_(d/D), d_G_ = k_on_G_max_(d/D), w_0_ = C_b_d_G_/(C_b_ + K_d_+K_d_d_G_), w_1_ = k_on_(C_b_+K_d_)^2^/(C_b_+K_d_+ K_d_d_G_).

<u>Desorption, with G(t=t_off_) = G_off_ and C_b_= 0 for t ≥ t_off_:</u>

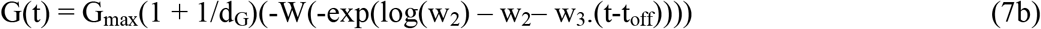

with w_2_ = (G_off_/G_max_).(1+1/d_G_), w_3_ = k_on_ K_d_/(1+d_G_).

Equations (7) are the binding equations of the 5-parameter (C_b_, G_max_, k_on_, K_d_, d/D) unstirred layer model. The Lambert W-function, also called ProductLog-function, in these equations is the principal solution W(x) of the equation e^w(x)^.W(x) = x. It is a standard mathematical function [22], implemented in programs like Wolfram Mathematica, Matlab, Maxima and a spreadsheet program like Origin.

From x = W(x)exp(W(x)) one finds dW(x)/dx = W(x)/(xW(x)+x) and using this relation one may verify that G(t), as given by equations (7), satisfies equation (6).

Because W(exp(w_0_ – w_1_.t)) → 0 for t → ∞, G(t) in equation (7a) will eventually reach the equilibrium value G_eq_. One may also verify that for t = 0 the second term in equation (7a) equals G_eq_, i.e. that G(t=0) = 0, but this requires extensive calculation and is much easier obtained by using a computer algebra program like Wolfram Mathematica.

Using W(exp(x)) requires attention because, for increasing values of x, the argument of W may become so large as to cause numerical overflow. This was prevented by introducing a new function Wexp(x) = W(exp(x)) that can be evaluated for all possible x.

### The new 4 -parameter “Surface interaction” binding model (Surfint)

We propose an adaptation of the Langmuir model in which incoming molecules may interact with adsorbed molecules and the adsorption rate will be influenced by the surface concentration G(t) to a degree G(t)/G_eq_. Without such molecular interaction we find from equation (2) for the Langmuir model that the adsorption rate v(t) diminishes at a rate proportional to itself: dv(t)/dt = -k_on_(C_b_+K_d_)v(t) and for the Surfint model we thus assume:

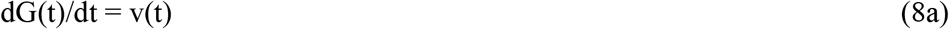

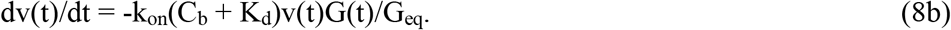

For the desorption rate v(t), with initial surface coverage G_eq_, we similarly assume that it decreases at a rate proportional to v(t) itself and the loss of protein (G_eq_ – G(t))/G_eq_:

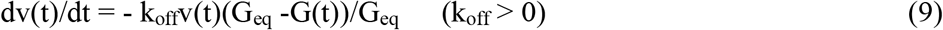

Equations (8) and (9) have the following solutions:

<u>Adsorption, with G(t=0)=0 and for t ≥ 0 a bulk concentration C_b_ of adsorbing protein:</u>

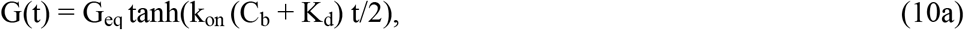

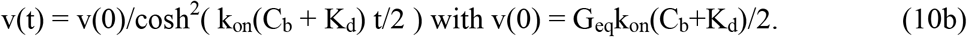

<u>Desorption, with G(t=t_off_) = G_eq_ and C_b_= 0 for t ≥ t_off_:</u>

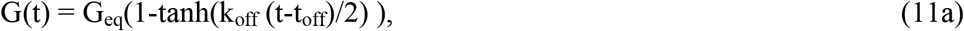

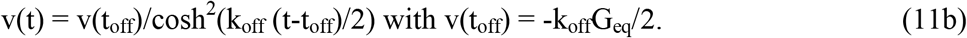

Equations (10) and (11) present the 4-parameter (C_b_, k_on_, k_off_, G_eq_) Surfint model. In contrast to the Langmuir model (for small non-interacting molecules located on large binding sites) the Surfint model has no G_max_ parameter, because for increasing surface concentration molecular interaction may change the packing mode of the adsorbed layer on the surface.

### A Random Sequential Adsorption (RSA) model with molecular surface interaction

A computer simulation of randomly adsorbing hard disks showed that maximally 54.7 % of total surface area is covered, leaving empty spots too small to fit a disk [23]. Mechanisms for closer packing have been suggested, such as “rolling” [24] or electrostatic deflection [25] of incoming molecules to neighbouring positions of adsorbed disks. At the buffer pH 7.4 in the present study, thrombin with pI = 7.1 [26,27] and prothrombin with pI = 4.6 [27,28] are both negatively charged, but for positively charged molecules a similar effect would occur.

We simulated molecular packing on a rectangular surface with a size of LxL (L=200) with periodic boundary conditions, i.e. what disappears on the right/top side returns on the left/bottom side and vice versa. The surface is hit by one sphere (with radius r = 1) per time unit (event A) at a random location. The total number of hits thus indicates time, and we considered the following possibilities:

1. The sphere hits an empty area and adsorbs on the surface (event B).
2. The sphere hits a single already adsorbed sphere (event C); then we return to event A with the nearest spot on the surface touching the adsorbed sphere as its location.
3. The sphere would hit 2 already adsorbed spheres (event D); then we return to event A with the nearest spot on the surface touching both adsorbed spheres as its location.
4. The sphere would hit 3 already adsorbed spheres. 4a When the 3 spheres form an acute triangle (event E): the sphere cannot be adsorbed. 4b When the 3 spheres form an obtuse triangle (event F): its new location touches the 2 most separated spheres in the triangle and we go back to event A.
5. When event F happens twice for the same sphere, the sphere cannot be adsorbed.

As shown in Fig.1, this procedure simulates spheres falling on the surface at a fixed time interval, the time unit, being disturbed in their fall if they hit previously absorbed spheres. If they can reach the nearest spot on the surface they will adsorb on it, otherwise they disappear. As shown in Fig.2, the formation of aggregates soon becomes dominant and, unexpectedly, we found equal final numbers of aggregates of 2 and 3 spheres. Desorption was added by giving each adsorbed sphere a fixed chance on disappearing per time unit.

**Fig.1.**
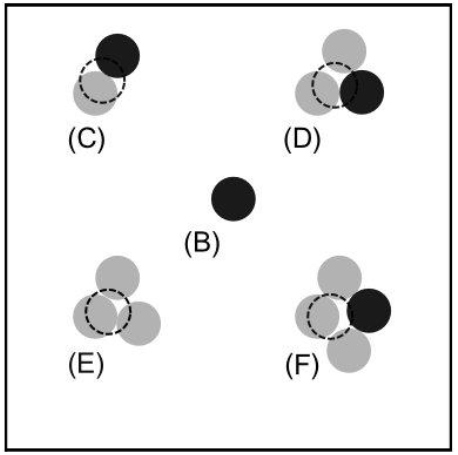
Incoming and final positions of spheres in the RSA model. Previously adsorbed spheres are shown in gray. Newly arriving spheres are shown as dashed, and their final positions in black. B to F indicate events as mentioned in the text.

**Fig.2.**
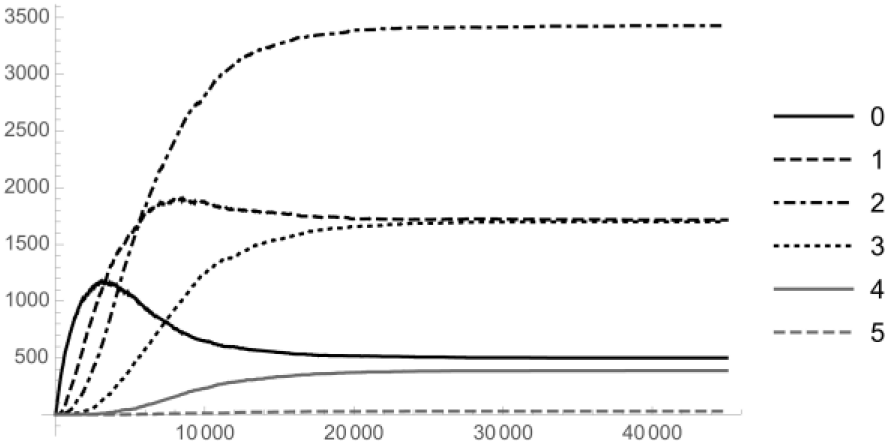
Time course of the numbers of isolated and aggregated spheres in the RSA model. Numbers of adsorbed sphere aggregates after random dropping of 45,000 spheres. Figures, added to the line types in the insert, indicate numbers of neighbouring spheres.

For small r/L ratio, sorption behaviour is independent of the size of r or L and this was already obained for r/L = 0.005 in which case 50,000 drops were sufficient to obtain an equilbrium state. Net desorption was studied by interrupting the arrival of new spheres and continuing desorption for another 25,000 time units. Such a single simulation, executed with Wolfram Mathematica 11 on a single core of a 3.2 GHz Intel Core i5, typically took 53 min.

### Ellipsometry

Unless specified differently, reagents were from Sigma-Aldrich (Zwijndrecht, The Netherlands). “Water” was de-ionized water (Milli-Q3, Millipore, Etten Leur, The Netherlands) and “buffer” was 20 mmol/L Hepes buffer, with physiological pH 7.4, and containing 140 mmol/L NaCl. Human thrombin and prothrombin were obtained from Synapse B.V., (Maastricht, The Netherlands) and adsorption is given in picomoles.cm^−2^, using molecular weights of M = 36000 for thrombin and M = 72000 for prothombin.

An automated Rudolph & Sons ellipsometer, type 4304-200E, was used as described [29]. Silica (SiO_2_)-coated silicon wafers (Wacker Chemitronic, n-type, phosphorus doped) were obtained from Aurel GmbH (Landsberg, Germany) and cut into slides of 4.0 × 0.8 cm. Protein binding on such pre-treated slides was measured at room temperature (21°C) in a glass cuvette with 3 ml of buffer containing a rotating magnetic stirrer (1150 rpm). Flow conditions in this system have been described [20] and total adsorbing surface area on the slides was 0.64 cm^2^. Light from a He/Ne-laser (Spectra Physics, ***λ=*** 632.8 nm) passes a polarizer P, hits the reflecting slide at an angle of incidence of 68 degrees, and then passes a second polarizer A. Both polarizers are rotated such that the final light intensity, measured by a photodiode, becomes minimal (null ellipsometry). From the positions of P and A adsorbed protein mass could be calculated every 10-14 sec as 85 ng/cm^2^ per degree change of the polarizer with a precision of about 1 ng/cm^2^ [30]. Desorption was measured after rapid (2-3 s) flushing of the cuvette with buffer. Times of onset of adsorption and desorption were obtained by linear extrapolation of the first two altered P values to baseline.

### Pre-treatment of ellipsometer cuvettes

The measured amounts of adsorbed proteins on the pretreated silicon slides were small enough to assume constant buffer concentrations C_b_ but this required prevention of aspecific protein adsorption on the inner cuvette walls by filling the cuvettes with organo-polysiloxane in heptane (Sigmacote) for 5 min with subsequenr drying in a stream of hot air. Excess siloxane was then removed by rubbing with cotton swabs and detergent (Sparkleen, Fisher Scientific Co, Pittsburg, Canada) followed by flushing with hot tap water. Finally, cuvettes were filled for 20 min with a solution of 1 g.L^−1^ of bovine serum albumin (BSA, ICN Biomedicals Inc., USA) at room temperature and then flushed with water. This procedure coats the inner cuvette walls with a layer of (denatured) BSA that prevents protein binding, as checked by ellipsometry on cuvette glass. Adsorption was studied in the Hepes/NaCl buffer.

### Binding of thrombin on heparin-coated slides

Slides were thoroughly cleaned by the following procedure. After flushing with detergent and hot tap water, the dried slides were kept for 10 s into a 4% hydrogen fluoride solution, flushed with water, dried, and kept for 1.5 h in 30% chromic sulfuric acid (80 g K_2_Cr_2_O_7_ with 250 mL H_2_SO_4_ and 750 mL water) (Merck, Darmstadt, Germany) at 80°C, Finally the slides were flushed with water, dried, covered with 20% carboxysilane (ABCR GmbH & Co, Karlsruhe, Germany) in 0.2 M sodiumactate (pH 4) and kept in an oven for 1 h at 110 °C. Hydrazine was coupled covalently to these slides at room temperature, using the standard method with N-hydroxy-sulfosuccinimide (NHS) and ethyl-dimethyl-cabodiimide (EDC) [31]. Such treatment changed the carboxy groups on the slides to hydrazide groups that bind spontaneously to aldehyde groups.

Mucosal porcine heparin (UFH) was obtained from Diosynth, Oss, The Netherlands, and was fractionated by gel filtration on a Superdex 75 Prep Grade column. Pooled fractions with molecular weights between M = 10,000 and M = 15,000 were used and characterized with respect to their physico-chemical properties [32,33]. At pH = 7.4 the heparin polysaccharide chains on the silica surface will carry a strong negative charge, because all three -OSO_3_^−^, - NHSO3^−^ and -COO – acid groups will be fully dissociated [34]. To produce aldehyde groups in this heparin, UFH was oxidized by overnight incubation of 5 mg/mL of heparin with 0.165 mg/mL sodium meta periodate in 50 mmol/L fosfate buffer (pH 6.9), at room temperature and in the dark. Finally, 67 ± 8 ng/cm^2^ of heparin (mean ± SD, n = 6) was coupled to the slide by adding 0.25 mg/mL of oxidized heparin to the hydrazine-activated slide in 0.4 mol/L acetate buffer (pH = 4.6) and overnight incubation at room temperature. Without induction of aldehyde groups, no thrombin adsorbed.

### Binding of prothrombin on supported phospholipid vesicles

Small unilamellar vesicles were prepared as described [29] by sonication of 80 mol% dioleoyl-phosphatidylserine (DOPS) and 20 mol% dioleoyl-phosphatidyl-choline (DOPC), both from Avanti Polar Lipids (Alabaster, USA). At the physiological pH = 7.4 used, the DOPS/DOPC vesicles are negatively charged because the phosphate (pI = 1-2) as well as the carboxylate (pI ≈ 4) group will be dissociated [35]. Phospholipid vesicles (20 μmol of lipid/L) were adsorbed on the silica slide surfaces, treated as described in the preceding section, in buffer with 2 mmol/L EDTA and pH 4.5. Such low pH (close to the isoelectric point of DOPS) was required for rapid vesicle adsorption to ± 680 ng.cm^−2^. For the vesicle diameter of 25 nm [36] and bilayer lipid mass of 400 ng.cm^−2^ [29], this was about 50% packing density.

The vesicle layers were then conditioned by alternate flushing with buffer with 10 mmol/L CaCl2 or 2 mmol/L EDTA, causing vesicle loss until a stable baseline of sparsely distributed vesicles was obtained of about 160 ng.cm^−2^. (This behaviour contrasts with adsorbed 20 mol% DOPS/80 mol% DOPC vesicles, fusing into bilayers which do not desorb upon flushing with buffer [29].) Binding of prothrombin to these vesicle is calcium-dependent and was studied in buffer containing 5 mmol/L CaCl_2_.Without phospholipid, no prothrombin adsorbed.

Prothrombin binding is reversible [37] but the high binding affinity makes desorption very slow. As a result, only partial desorption could be measured. Also, few C_b_ values were used because for C_b_> 40 pmol.cm^−3^ the fast initial upstroke of G(t) contained few data points, and for C_b_ < 10 pmol.cm^−3^ adsorption lowered the buffer concentration C_b_. For the experiments shown, such lowering was less than 1%.

### Parameter fitting. Uncertainty and interdependence of obtained parameter values

The quality of obtained fits is expressed by the weighted L_2_-norm of the residue:

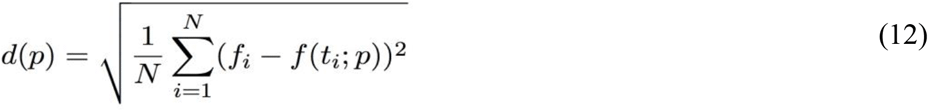

with f(t;p) the solution of the differential equation, p the unknown parameter vector and ***i*** a measured point in the curve. Equation (12) presents the average distance between measured and fitted surface concentrations, in pmol.cm^−2^. Minimization of d(p) was performed with Mathematica. [Wolfram Research, Inc., Mathematica, Version 10.2, Champaign, IL (2015).] Several numerical approaches produced similar results and the most efficient and robust minimization method, Levenberg-Marquardt iteration [38], was used for the results presented.

For normal distribution of error, parameter confidence limits are determined from Fisher’s F-distribution as elliptical areas, indicating uncertainty and interdependence [39]. However, because of systematic differences between measured and fitted values (see Fig.5), error was not stochasticly independent and exact confidence intervals cannot be calculated. Therefore, parameter uncertainty and interdependency, shown in Fig.6, show the regions where d(p) remains within 110%, 120%, 150% and 200% of the minimal value d(p_0_).

## Results

Fig.3 shows the results of a 5 times repeated RSA experiment, resulting in total adsorption of 7756, 7741, 7736, 7762 and 7736 spheres, i.e. 60.92, 60.80, 60.76, 60.96 and 60.76 percent coverage of total surface area. This is higher than the 54.7 % obtained for disks not adsorbing when hitting already adsorbed disks [23], but still far from the maximal value of 90.7%, as obtained for irreversibly adsorbed disks taking part in lateral diffusion [7]. The 5 RSA curves in Fig.3 coincided within line width with the function 7731.tanh(0.0001345 x).

**Fig.3.**
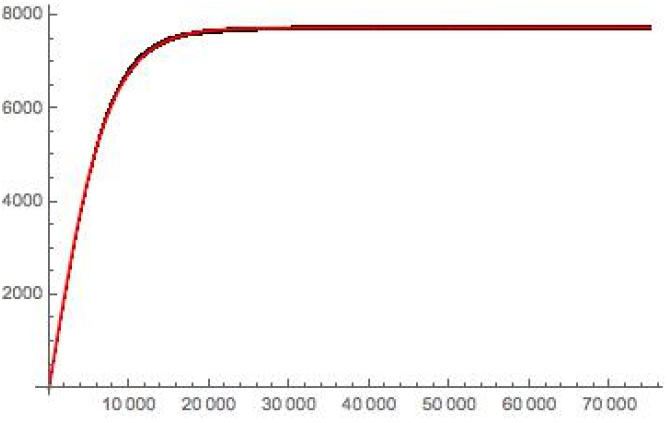
Numbers of adsorbed spheres in RSA simulations. Results of five RSA experiments with dropping of a total number of 76,000 spheres, with best-fitting tanh-function (see text). The six lines coincide within line width.

Fig.4 shows a similar experiment including desorption. At the arrival of each new sphere, i.e. after each time unit, all adsorbed spheres had a fixed chance on desorption, 0.003 % for the upper curve and 0.03 % for the lower curve, causing exponential desorption. After dropping of 50,000 spheres (t_off_=50,000), adsorption was stopped and net desorption is shown for another 25,000 time units. Again adsorption closely followed 7134.tanh(0.000136 t) (upper curve) and 3001.tanh(0.000252 t) (lower curve) (t< t_off_) and desorption the exponential functions 7134.exp(−0.0000293(t-t_off_)) and 3001.exp(−0.000296(t-t_off_)), (t > t_off_), respectively. The 6 curves in Fig.4 again fall close together but comparison with Fig.3 shows that addition of desorption slightly increased scatter. It is concluded that, RSA simulation confirmed the Surfint model, with tanh sorption functions, also in the presence of desorption.

**Fig.4.**
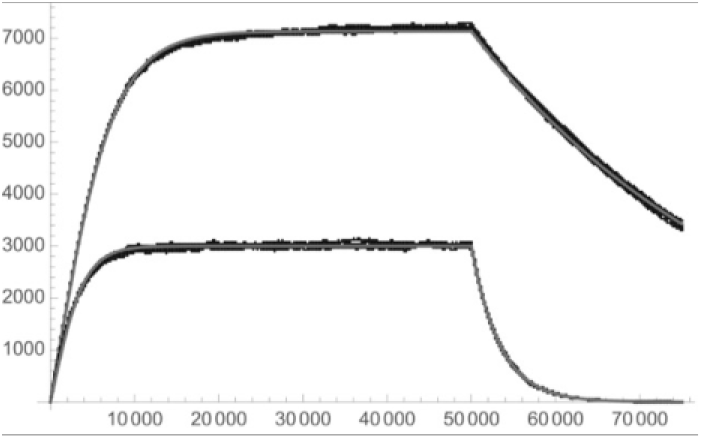
RSA simulations with desorption. During and after dropping of 50,000 spheres, each adsorbed sphere had a chance on desorption (see text). Best-fit tanh adsorption functions and exponential desorption functions are shown for two different desorption constants (see text).

As shown in Fig.5 and Table1, thrombin adsorptions fitted the Surfint model well, with a mean residue of 0.0258 pmol.cm^−2^ or 0.93 ng.cm^−2^, i.e.within ellipsometric accuracy of 1 ng.cm^−2^, and no systematic deviations between experimental and fitted curves. Surfint parameters were well-defined, as shown in Fig.6, and k_on_ values in Table 1 decreased by a factor of 5 for increasing surface coverage Geq, confirming the influence of surface coverage on adsorption kinetics. The Langmuir and Unstir models fitted thrombin sorption worse, with mean residues of 0.0523 and 0.0416 pmol.cm^−2^, respectively, and systematic deviations.

**Fig.5.**
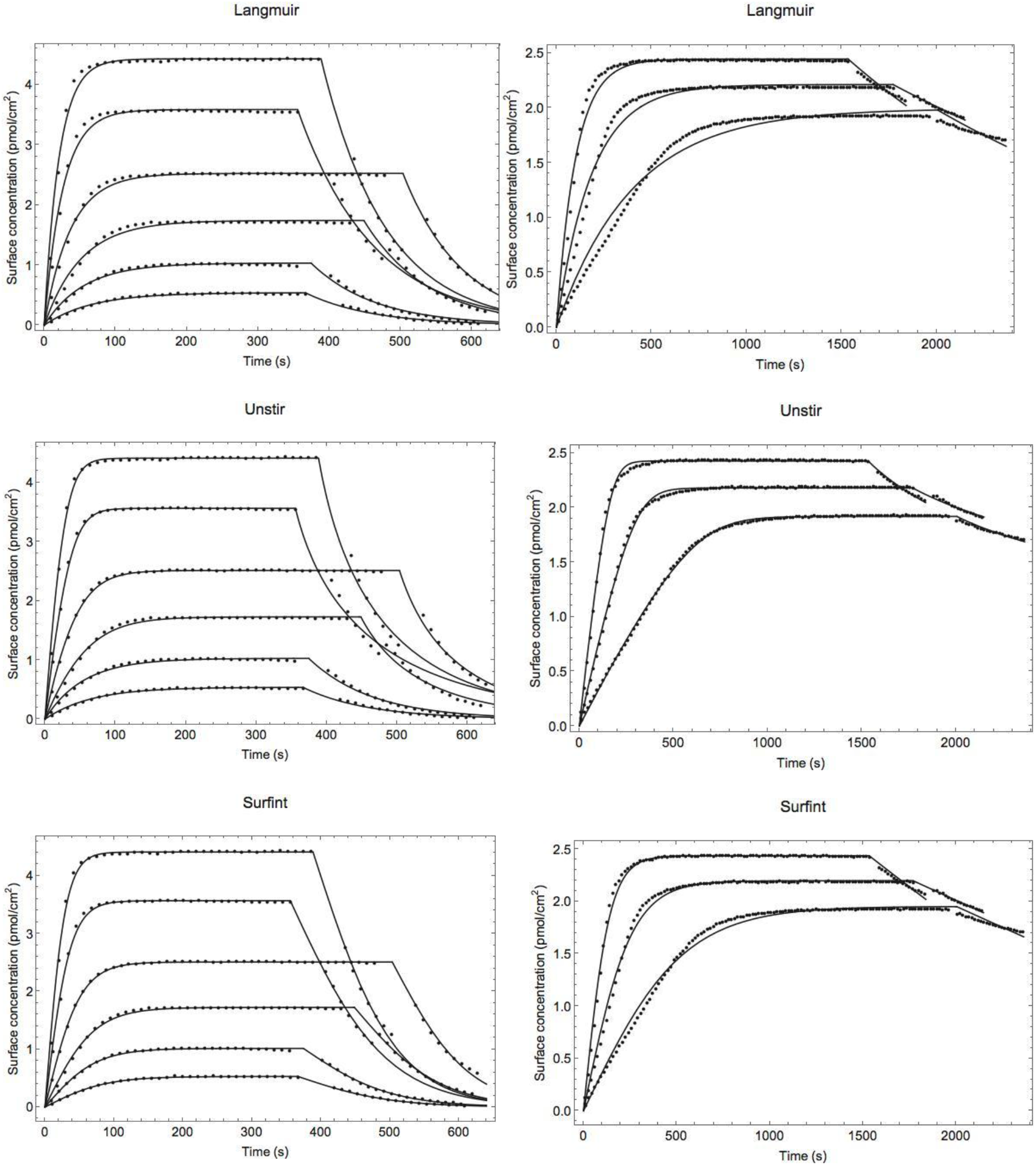
Obtained fits for the three models. Fits of the Langmuir, Unstir and Surfint models for the binding of thrombin on heparin (left figures) and the binding of prothrombin on 80% DOPS/20% DOPC phospholipid vesicles (right figures).

**Fig.6.**
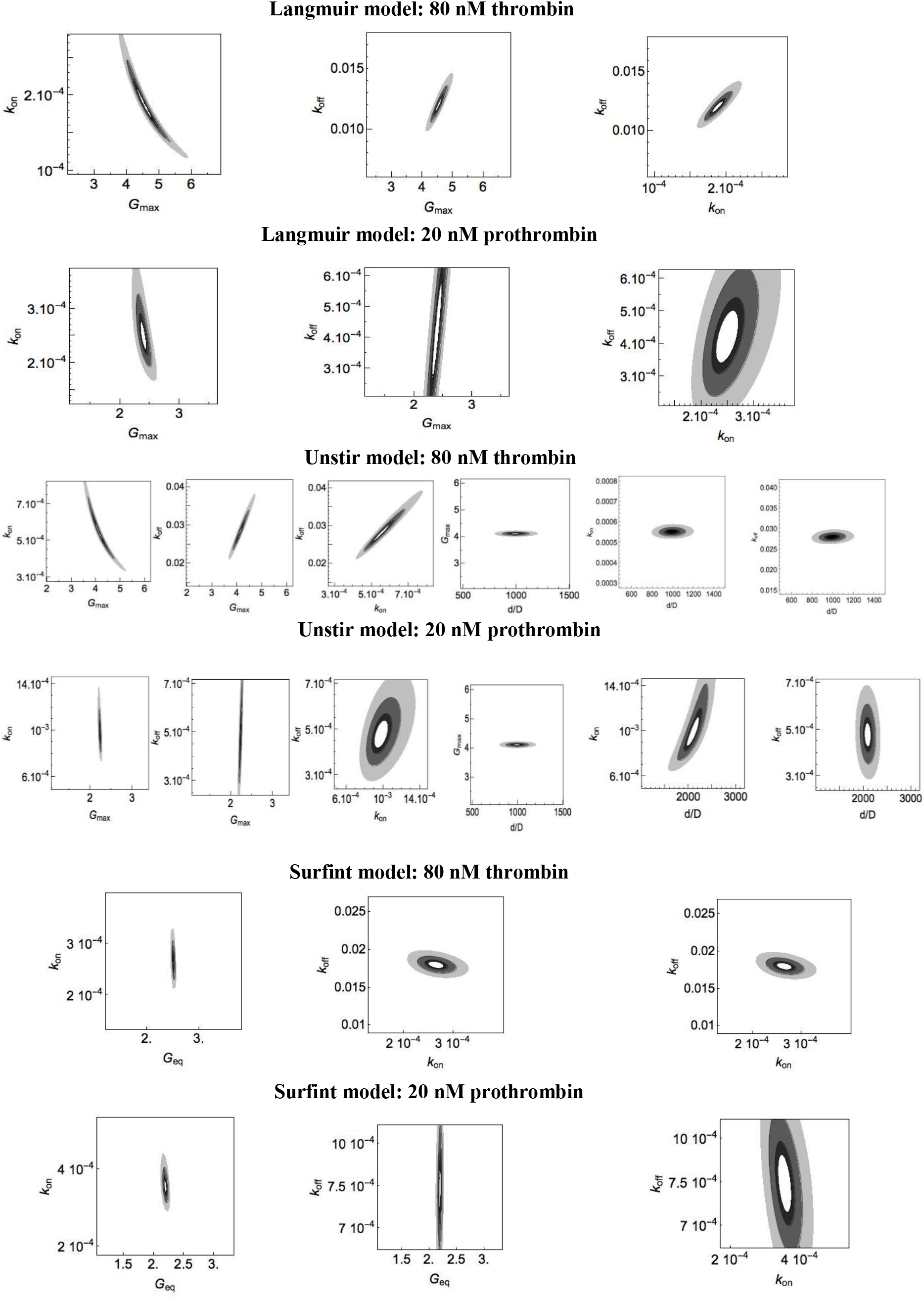
Parameter determination and parameter interdependence. Indicated regions present residue values of 110% (white), 120% (black), 150% (dark grey) and 200% (light gray) of the minimal residue value d(p_0_).

**Table 1:**
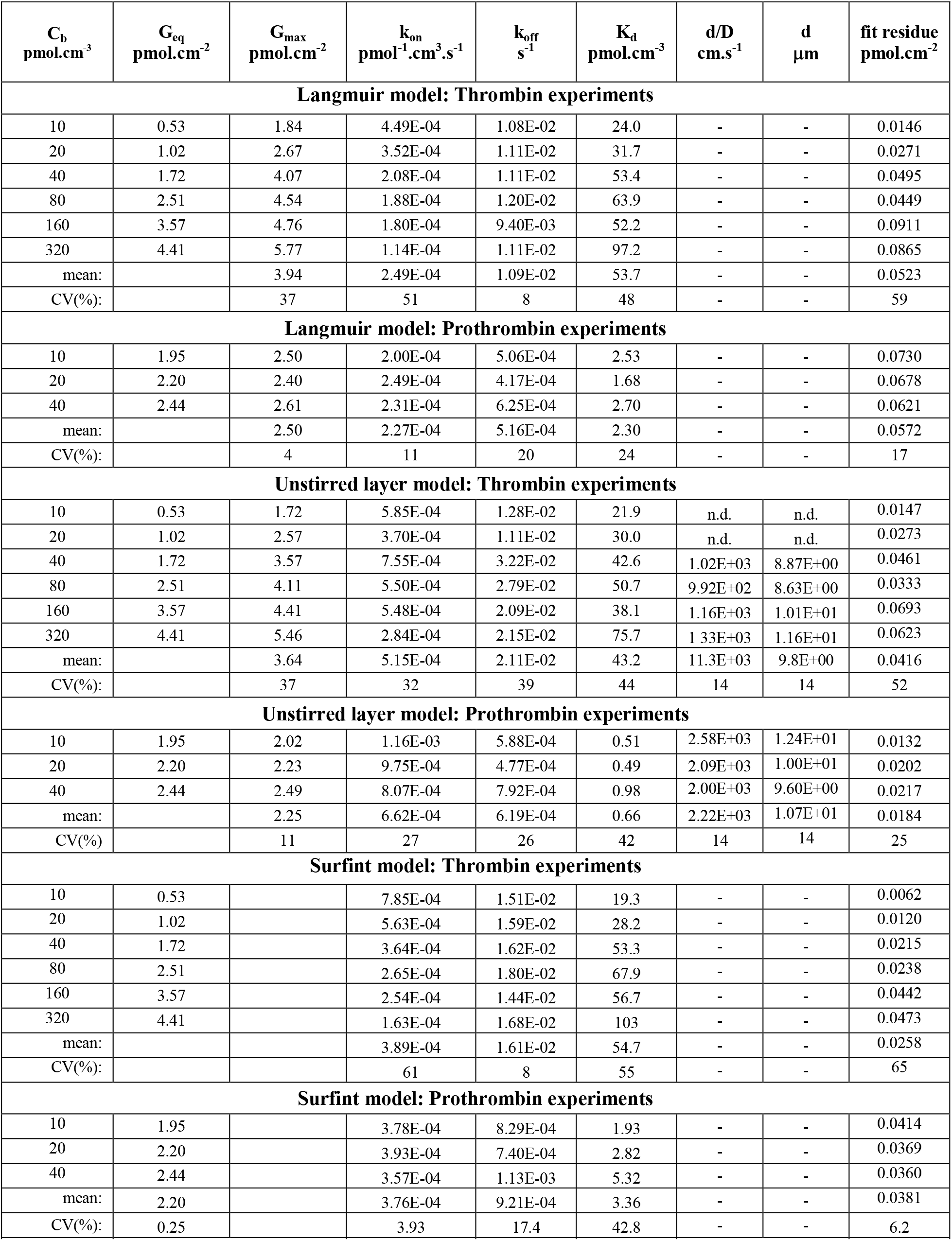
Obtained binding parameters.

| $C_b$<br>pmol.cm<sup>-3</sup> | $G_{eq}$<br>pmol.cm<sup>-2</sup> | $G_{max}$<br>pmol.cm<sup>-2</sup> | $k_{on}$<br>pmol<sup>-1</sup>.cm<sup>3</sup>.s<sup>-1</sup> | $k_{off}$<br>s<sup>-1</sup> | $K_d$<br>pmol.cm<sup>-3</sup> | d/D<br>cm.s<sup>-1</sup> | d<br>μm | fit residue<br>pmol.cm<sup>-2</sup> |
| --- | --- | --- | --- | --- | --- | --- | --- | --- |
| **Langmuir model: Thrombin experiments** |  |  |  |  |  |  |  |  |
| 10 | 0.53 | 1.84 | 4.49E-04 | 1.08E-02 | 24.0 | - | - | 0.0146 |
| 20 | 1.02 | 2.67 | 3.52E-04 | 1.11E-02 | 31.7 | - | - | 0.0271 |
| 40 | 1.72 | 4.07 | 2.08E-04 | 1.11E-02 | 53.4 | - | - | 0.0495 |
| 80 | 2.51 | 4.54 | 1.88E-04 | 1.20E-02 | 63.9 | - | - | 0.0449 |
| 160 | 3.57 | 4.76 | 1.80E-04 | 9.40E-03 | 52.2 | - | - | 0.0911 |
| 320 | 4.41 | 5.77 | 1.14E-04 | 1.11E-02 | 97.2 | - | - | 0.0865 |
| mean: |  | 3.94 | 2.49E-04 | 1.09E-02 | 53.7 | - | - | 0.0523 |
| CV(%): |  | 37 | 51 | 8 | 48 | - | - | 59 |
| **Langmuir model: Prothrombin experiments** |  |  |  |  |  |  |  |  |
| 10 | 1.95 | 2.50 | 2.00E-04 | 5.06E-04 | 2.53 | - | - | 0.0730 |
| 20 | 2.20 | 2.40 | 2.49E-04 | 4.17E-04 | 1.68 | - | - | 0.0678 |
| 40 | 2.44 | 2.61 | 2.31E-04 | 6.25E-04 | 2.70 | - | - | 0.0621 |
| mean: |  | 2.50 | 2.27E-04 | 5.16E-04 | 2.30 | - | - | 0.0572 |
| CV(%): |  | 4 | 11 | 20 | 24 | - | - | 17 |
| **Unstirred layer model: Thrombin experiments** |  |  |  |  |  |  |  |  |
| 10 | 0.53 | 1.72 | 5.85E-04 | 1.28E-02 | 21.9 | n.d. | n.d. | 0.0147 |
| 20 | 1.02 | 2.57 | 3.70E-04 | 1.11E-02 | 30.0 | n.d. | n.d. | 0.0273 |
| 40 | 1.72 | 3.57 | 7.55E-04 | 3.22E-02 | 42.6 | 1.02E+03 | 8.87E+00 | 0.0461 |
| 80 | 2.51 | 4.11 | 5.50E-04 | 2.79E-02 | 50.7 | 9.92E+02 | 8.63E+00 | 0.0333 |
| 160 | 3.57 | 4.41 | 5.48E-04 | 2.09E-02 | 38.1 | 1.16E+03 | 1.01E+01 | 0.0693 |
| 320 | 4.41 | 5.46 | 2.84E-04 | 2.15E-02 | 75.7 | 1.33E+03 | 1.16E+01 | 0.0623 |
| mean: |  | 3.64 | 5.15E-04 | 2.11E-02 | 43.2 | 11.3E+03 | 9.8E+00 | 0.0416 |
| CV(%): |  | 37 | 32 | 39 | 44 | 14 | 14 | 52 |
| **Unstirred layer model: Prothrombin experiments** |  |  |  |  |  |  |  |  |
| 10 | 1.95 | 2.02 | 1.16E-03 | 5.88E-04 | 0.51 | 2.58E+03 | 1.24E+01 | 0.0132 |
| 20 | 2.20 | 2.23 | 9.75E-04 | 4.77E-04 | 0.49 | 2.09E+03 | 1.00E+01 | 0.0202 |
| 40 | 2.44 | 2.49 | 8.07E-04 | 7.92E-04 | 0.98 | 2.00E+03 | 9.60E+00 | 0.0217 |
| mean: |  | 2.25 | 6.62E-04 | 6.19E-04 | 0.66 | 2.22E+03 | 1.07E+01 | 0.0184 |
| CV(%): |  | 11 | 27 | 26 | 42 | 14 | 14 | 25 |
| **Surfint model: Thrombin experiments** |  |  |  |  |  |  |  |  |
| 10 | 0.53 |  | 7.85E-04 | 1.51E-02 | 19.3 | - | - | 0.0062 |
| 20 | 1.02 |  | 5.63E-04 | 1.59E-02 | 28.2 | - | - | 0.0120 |
| 40 | 1.72 |  | 3.64E-04 | 1.62E-02 | 53.3 | - | - | 0.0215 |
| 80 | 2.51 |  | 2.65E-04 | 1.80E-02 | 67.9 | - | - | 0.0238 |
| 160 | 3.57 |  | 2.54E-04 | 1.44E-02 | 56.7 | - | - | 0.0442 |
| 320 | 4.41 |  | 1.63E-04 | 1.68E-02 | 103 | - | - | 0.0473 |
| mean: |  |  | 3.89E-04 | 1.61E-02 | 54.7 | - | - | 0.0258 |
| CV(%): |  |  | 61 | 8 | 55 | - | - | 65 |
| **Surfint model: Prothrombin experiments** |  |  |  |  |  |  |  |  |
| 10 | 1.95 |  | 3.78E-04 | 8.29E-04 | 1.93 | - | - | 0.0414 |
| 20 | 2.20 |  | 3.93E-04 | 7.40E-04 | 2.82 | - | - | 0.0369 |
| 40 | 2.44 |  | 3.57E-04 | 1.13E-03 | 5.32 | - | - | 0.0360 |
| mean: | 2.20 |  | 3.76E-04 | 9.21E-04 | 3.36 | - | - | 0.0381 |
| CV(%): | 0.25 |  | 3.93 | 17.4 | 42.8 | - | - | 6.2 |

For the Langmuir model Table 1 shows that k_on_ decreased by a factor of 4 for increasing concentrations of thrombin, which is not compatible with this model. It is concluded that thrombin adsorption was adequately described by the Surfint model but not by the Langmuir and Unstir models.

For the high-affinity binding of prothrombin, diffusion-limitation became important as shown by the well defined parameter d/D in Fig.6 and by the low residue of 0.018 pmol.cm^−2^, or 1.3 ng.cm^−2^, for the Unstir model in Table 1. In this case, the Surfint and Langmuir models, not incorporating diffusion, both showed systematic deviations in Fig.5, and larger mean residues of 0.038 and 0.057 pmol.cm^−2^. As further shown in Fig.6, most parameters are well-determined, except koff for prothrombin binding, because its slow desorption could only be partially measured. Using values of D = 8.76×10^−7^ cm^2^.s^−1^ for thrombin [40] and D = 4.8×10^−7^ cm^2^.s^−1^ for prothrombin [41], the corresponding unstirred layer thickness of 9.8 μm and 10.7 μm in Table 1 is within the range of reported values of 2.5-12 μm [20,42,43].

## Discussion

The importance of surface-related processes in biology is illustrated by the fact that 26% of all human genes code for membrane-bound proteins [44] and about half of all intracellular proteins are membrane-associated [45]. In addition, many plasma proteins need binding to glycogen, collagen or phospholipids to become functional. Such binding induces reactive protein conformations but also enhances the probability of meeting reaction partners by so called “reduction of dimensionality” [46], i.e. restriction of molecular movement to a (heparin) chain or a (vesicle) surface, instead of random movement in solution.

This is also true for the proteins used in the present study. To be activated to thrombin, prothrombin requires binding of its phosphorylated *γ*-glutamin residues to PS-containing membranes [47]. For the binding of thrombin to heparin, at least 12 sugar residues in the heparin chain are required, allowing it to meet its physiological inhibitor anti-thrombin III, bound on the same heparin chain [33,48]. Similar examples have been presented for protein transport along DNA chains [45].

Such (previously unknown) binding requirements may explain why reported K_d_ values for the binding of thrombin to heparins varied between 1 to 1000 nM [48] but the presently measured value of 53.7 nM in Table 1 for the Langmuir model is in agreement with the K_d_ value of 61 nM reported for the binding of thrombin to biotinylated heparin immobilized on streptavidin-coated sensors [49].

For the binding of prothrombin to phospholipid vesicles, most studies in the literature used lower (more physiological) PS/PC ratios [5,20,30,47]. But the K_d_ value of 2.3 nM shown in Table 1 for the Langmuir model is in the range of the values of 4.5 nM, 7 nM and 8 nM reported for prothrombin binding on stacked 100% DOPS mulilayers [37], 1:2 POPC/POPS monolayers [50] and pure brain PS [51], repectively. These studies also used the Langmuir model but, for such high-affìnity, binding becomes transport-limited and the Unstir model is more appropriate. This is illustrated in the present study by the difference between mean K_d_ values, obtained for the Langmuir model (2.3 nM) and for the Unstir model (0.66 nM).

The frequent use of the Langmuir model is based on its simplicity, with fixed constants k_on_, k_off_ and G_max_, allowing simultaneous fitting of a series of sorption curves with different buffer concentrations C_b_. As shown in Table 1, however, the k_on_ values as obtained for the Langmuir model were not constant but decreased by a factor of 4 for increasing C_b_. This also indicates an effect of surface coverage on k_on_. Also, the mean residues of 0.052 and 0.057 pmol.cm^−3^ in Table 1, for separate fits of thrombin and prothrombin adsorptions, increased to 0.073 and 0.097 pmol.cm^−^ for simultaneous fits (results not shown), indicating that the assumption of constant sorption constants is not valid.

In contrast to the adsorption constants, desorption constants were rather constant for all three models, indicating that desorption was not dependent on surface coverage with protein. For the buffer pH of 7.4, the heparin and DOPS/DOPC surfaces are both strongly negatively charged and the additional negative charges of the protein molecules on the surface seem to have little influence on desorption. Once adsorbed, the interaction with the surface seems to dominate over interactions between adsorbed molecules.

We previously proposed [52] that the influence of surface coverage on sorption kinetics could explain so-called Vroman effects, i.e. time-dependent changes in the composition of adsorbed protein layers in contact with a multi-protein solution. The present study shows that the influence of macromolecular surface coverage on sorption kinetics can be formulated in a more general form by using the Surfint model. A possible underlying molecular mechanism for such influence was presented in the RSA model, showing that interaction between incoming and adsorbed molecules may cause tanh adsorption kinetics, with a longer-lasting initial linear upstroke than the exponential function of the Langmuir model. In the RSA model, the proposed mechanisms - electrostatic or steric deflection of incoming molecules to positions adjacent to adsorbed molecules - extends the lineair adsorption phase because it makes the adsorbed molecules invisible for further adsorption until the number of 3-particle aggregates becomes significant. The effect of surface coverage on the adsorption rate v(t) can also be related to surface entropy S_surf_, which according to Boltzmann’s relation S_surf_ = k ln(W), with k the Boltzmann constant and W the number of particle position permutations, remains low in clustered configurations [25].

## Conclusion

The present study shows shortcomings of the well known Langmuir binding model for description of macromolecular binding because the model does not include interaction between incoming and adsorbed molecules. Also, the model does not include transport limitation, as will occur for high-affinity binding. To solve the first problem, we introduced the new Surfint model and demonstrated its better fitting of low-affinity binding of thrombin on heparinized slides. For transport-limited adsorption molecular interaction no longer influences sorption kinetics and binding can be best described by the Unstir model. This is shown for the high-affinity binding of prothrombin. Both for the Surfint and for the Unstir model we present the exact analytical solution for the differential equations that describe the model.

